# Methylmalonic acid induces metabolic abnormalities and exhaustion in CD8^+^ T cells to suppress anti-tumor immunity

**DOI:** 10.1101/2024.03.03.583124

**Authors:** Joanne D. Tejero, Rebecca S. Hesterberg, Stanislav Drapela, Didem Ilter, Devesh Raizada, Felicia Lazure, Hossein Kashfi, Min Liu, Juan Fernández-García, John M. Asara, Sarah-Maria Fendt, John L. Cleveland, Ana P. Gomes

## Abstract

Systemic levels of methylmalonic acid (MMA), a byproduct of propionate metabolism, increase with age and MMA promotes tumor progression via its direct effects in tumor cells. However, the tumorigenic role of MMA in modulating the tumor ecosystem remains to be investigated. The proliferation and function of CD8^+^ T cells, key anti-tumor immune cells, declines with age and in conditions of vitamin B12 deficiency, the two most well-established conditions that lead to increased systemic levels of MMA. Thus, we hypothesized that increased circulatory levels of MMA leads to suppression of CD8^+^ T cell immunity. Treatment of primary CD8^+^ T cells with MMA induced a dysfunctional phenotype characterized by a robust immunosuppressive transcriptional reprogramming and marked increases in the expression of the exhaustion regulator, TOX. Accordingly, MMA treatment upregulated exhaustion markers in CD8^+^ T cells and decreased their effector functions, which drove the suppression of anti-tumor immunity *in vitro* and *in vivo*. Mechanistically, MMA-induced CD8^+^ T cell exhaustion was associated with a suppression of NADH-regenerating reactions in the TCA cycle and concomitant defects in mitochondrial function. Thus, MMA has immunomodulatory roles, thereby highlighting MMA as an important link between aging, immune dysfunction, and cancer.

## Introduction

Methylmalonic acid (MMA) is a metabolite produced as a byproduct of propionate metabolism, a mitochondrial pathway that catabolizes branched chain amino acids, odd chain fatty acids, and cholesterol to fuel the tricarboxylic acid (TCA) cycle. MMA levels are generally low under normal physiological conditions, yet its levels rise in pathological conditions. Chief amongst these are conditions of vitamin B12 deficiency [1, 2]. Importantly, multiple studies have shown that increases in circulatory MMA levels are a feature of aging [3–6]. Considering that aging is a major risk factor for development of cancer [7, 8], the relationship between increased MMA levels and cancer has been explored. Increased MMA in circulation has been shown to promote tumor progression into metastatic disease through direct induction of aggressive features in the tumor cells [6]. On the other hand, MMA production can also be hijacked by non-age-related aggressive tumors to enable their cell autonomous acquisition of pro-metastatic properties [9], and influences the fibroblast compartment of the tumor microenvironment (TME) by skewing the fibroblasts toward a cancer-associated fibroblast phenotype [10]. However, how MMA impacts the other components of the TME that affect tumorigenesis has not been investigated.

CD8^+^ T cells are adaptive immune cells that are an important component of the TME, functioning to curb tumorigenesis and tumor progression through their cytotoxic activity [11, 12]. Thus, it comes as no surprise that tumor cells utilize an array of strategies to disable CD8^+^ T cell-mediated cytotoxic activity by driving their exhaustion [13], a state that normally occurs due to chronic antigen exposure and that functions as an important homeostatic response to prevent autoimmunity [14, 15]. A gradual decline in CD8^+^ T cell function is a central feature of the aging process, with CD8^+^ T cells exhibiting signs of exhaustion [16, 17]. CD8^+^ T cells also experience age-associated mitochondrial dysfunction, characterized by decreased mitochondrial oxidative phosphorylation and increased reactive oxygen species (ROS) [16, 17], features that are known to associate with, and in some cases induce, CD8^+^ T cell exhaustion [18–20]. Understanding the factors that drive CD8^+^ T cell exhaustion that occur with old age is instrumental for developing strategies to impair tumor initiation and progression.

Given that aging is associated with an increase in circulatory MMA as well as a decline in CD8^+^ T cell function, we investigated whether a cause-and-effect relationship exists between these two aging features and their potential consequences for tumorigenesis. Herein we show that such relationship exists, whereby MMA promotes CD8^+^ T cell exhaustion, driven by the defects in mitochondrial metabolism and the induction of the exhaustion master regulator TOX, which impairs anti-tumor responses and enables immune escape. Thus, MMA is a potent regulator of CD8^+^ T cell fate and function, and is an important biomarker of immune function that has the potential to be exploited as a predictor of immunotherapy responses.

## Materials and methods

A summary of all key materials and additional methodology is provided in the Supplementary Information.

### Mice

All animal studies were approved by the Institutional Animal Care and Use Committee (IACUC) of Moffitt Cancer Center and the University of South Florida. Male and female C57BL/6NJ mice or OT-I mice (a gift from Dr. Avram), aged 8 to 16 weeks, were used. Animals were group-housed and bred in standard cages in a specific pathogen-free animal facility. Animals were provided with unrestricted water and food (Teklad Irradiated Global 18% Protein Rodent Diet 2918; Envigo, Indianapolis, IN, USA). Animal husbandry was carried out by the vivarium technical staff. The room was maintained at 20-23°C with 30-70% humidity on a 12 hour light-dark cycle. Some animals were treated with MMA when indicated and the administration of MMA to mice was based on a previous study by Ribeiro, *et al.* [21]. Briefly, MMA (Sigma-Aldrich, St. Louis, MO, USA) was dissolved in 0.9% normal saline and buffered to pH 7.4 using sodium hydroxide. MMA (or saline) was intraperitoneally (i.p.) injected twice daily with an 8 hour interval between injections for 28 days. MMA doses were escalated every week: 0.76 μmol of MMA per gram of body weight at week 1, 1.01 μmol of MMA per gram of body weight at week 2, 1.86 μmol of MMA per gram of body weight at week 3, and 2.67 μmol of MMA per gram of body weight at week 4. Mice were weighed every other day. All injections were prepared such that the mice received 5 μL of solution per gram of body weight.

### Syngeneic transplantation of tumor cells

C57BL/6NJ male mice, aged 9 to 20 weeks, were randomly divided into two groups which received twice daily saline or MMA i.p. injections as described above. At day 4, tumor cells derived from KP^G12D^ mice (a genetically engineered lung cancer mouse model with LSL-Kras^G12D^;Trp53^flox/flox^) [22], which are tagged with a GFP-luciferase reporter (referred to as KP-Luciferase cells) were injected at 100,000 cells per 100 μL PBS via tail vein. At day 28, mice were euthanized, and spleen and lung tissues were collected for analysis by flow cytometry and histology.

### Mouse CD8^+^ T cell isolation and culture conditions

Single cell suspensions were prepared from the spleens and pooled lymph nodes of C57BL/6NJ or OT-I mice by mechanical dissociation through a 70 μm nylon cell strainer in separation buffer (SB: PBS + 3% FBS + 2 mM EDTA). Red blood cells were lysed using RBC lysis buffer (Biolegend, San Diego, CA, USA) for 5 minutes on ice. Cells were washed twice with SB, filtered through a 70 μm nylon cell strainer, and resuspended at concentration of 10^8^ cells/mL in SB. Naive CD8^+^ T cells were isolated by negative selection using the MojoSort Mouse CD8^+^ Naïve T cell Isolation Kit as per manufacturer’s instructions (BioLegend, >90% purity). Live cells were counted by trypan blue dye exclusion and resuspended at a concentration of 10^6^ cells/mL in complete T cell media (RPMI-1640 (Cytiva, Marlborough, MA, USA) with 10% FBS (Gibco, Billings, MT, USA), 1% penicillin/streptomycin (Cytiva), 50 μM β-mercaptoethanol (Sigma-Aldrich), 1 mM sodium pyruvate (Gibco), and 1% MEM nonessential amino acids (Gibco). Cells were seeded at a concentration of 100,000 cells/well in 96-well round-bottom plates (Sarstedt, Newton, NC, USA) that were coated overnight with 10 μg/mL of anti-mouse CD3ε antibody (Biolegend) and washed twice with PBS prior to seeding. On the same day of isolation, T cell activation was initiated using 0.5 μg/mL anti-mouse CD28 (BioLegend) and 10 ng/mL recombinant mouse interleukin-2 (IL-2) (BioLegend); additionally, MMA (Sigma-Aldrich), solubilized in molecular grade water, was added at a concentration of 5 mM for up to 72 hours at 37°C and 5% CO2.

### Cytotoxicity assay

Mouse CD8^+^ T cells treated with vehicle or 5 mM MMA for 3 days (as described above) were harvested, washed with PBS, and re-seeded with complete T cell media and 10 ng/mL IL-2 for an additional 2 days. KP-Luciferase cells were seeded at 3,000 cells/well in 96-well flat bottom plates and cultured with the conditioned media from activated CD8^+^ T cells for 3 days. At endpoint, propidium iodide was used to assess tumor cell death using Incucyte S3 Live-Cell Analysis System v2021A (Sartorius, Göttingen, Germany). Cell death was measured by calculating propidium iodide positive (red) object count normalized to confluency. In other experiments, CD8^+^ T cells were isolated from male OT-I mice (CD8^+^ OT-I cells) and activated in the presence of anti-CD3, anti-CD28, IL-2, and concurrently treated with vehicle or 5 mM MMA for 3 days in 96-well round bottom plates. These CD8^+^ OT-I cells were co-cultured with Lewis Lung Carcinoma cells that express ZsGreen or OVA-T2A-ZsGreen (LLC-OVA-ZsGreen or LLC-ZsGreen [23]) target cells at a 5:1 effector-to-target ratio for 24 hours in complete T cell media. Percent cytotoxicity was calculated using the following equation, where ZY^+^ represents zombie yellow positive (dead) cells: (ZY^+^LLC-OVA minus ZY^+^LLC without OVA) / (100 minus ZY^+^LLC without OVA) x 100.

### Flow cytometry

For assessment of cell surface markers, cells were washed at least once with PBS, and stained with the indicated antibodies (anti-CD3, anti-CD8, anti-CD25, anti-CD69, anti-PD-1, anti-CD38, anti-TIM-3, anti-LAG-3; detailed in Supplementary Information) for 20 to 30 minutes at 4°C in PBS prior to washing and analysis. For assessment of transcription factors, cells were first surface stained as described above and fixed (eBioscience Transcription Factor Staining Set; Thermo Fisher Scientific, Waltham, MA, USA) overnight at 4°C. The next day, cells were stained with transcription factor antibody (TOX; Thermo Fisher Scientific) in permeabilization buffer (eBioscience Transcription Factor Staining Set; Thermo Fisher Scientific) for 1 hour at 4°C prior to washing and analysis. For assessment of intracellular cytokines, cells were treated for 5 hours with a protein transport inhibitor containing brefeldin A (BD Biosciences, Franklin Lakes, NJ, USA), washed and fixed for 10 minutes at 4°C with fixation/permeabilization solution (BD Cytofix/Cytoperm kit) and washed with permeabilization buffer overnight before staining for 30 minutes with anti-TNF-α, anti-IFN-γ, and anti-granzyme B antibodies (detailed in Supplementary Information) prior to washing and analysis. For assessment of T cell proliferation, Cell Trace Violet (Thermo Fisher Scientific) dye was used according to manufacturer’s instructions, and the division index (the average number of cell divisions that a cell in the original population has undergone, including the undivided peak) was calculated. For assessment of reactive oxygen species, CM-H2DCFDA (Thermo Fisher Scientific) and MitoSOX Red (Thermo Fisher Scientific) were used according to manufacturer’s instructions. For assessment of viability, cells were stained with either propidium iodide (Sigma-Aldrich) or a fixable viability dye (Thermo Fisher Scientific). Data was collected using Attune NxT (Thermo Fisher Scientific) or BD FACSymphony (BD Biosciences) flow cytometers and analysis was performed on FlowJo v10.8.1 software (BD Biosciences).

### Statistical analysis

Statistical analyses were performed with GraphPad Prism v9.1.0 (GraphPad Software, Boston, MA, USA). Statistical significance was determined as a p-value ≤ 0.05. Student t-tests (paired or unpaired) were used to calculate the differences between groups. Data are represented as the mean ± SEM (standard error of the mean) of individual data points of at least three biological replicates.

## Results

### MMA impairs activation and effector functions of primary CD8^+^ T cells

A characteristic feature of CD8^+^ T cell dysfunction is the loss of proliferative ability upon activation [24, 25]. We first assessed how a pathologically relevant concentration of MMA (5 mM, a treatment concentration that mimics the intracellular concentration of MMA of cells exposed to aged serum [6]) affected CD8^+^ T cell proliferation. Evaluation of primary CD8^+^ T cells isolated from mice revealed that while MMA treatment had no effect on their viability (Supplementary Fig. 1A), it significantly decreased their proliferation, as determined by cell trace violet staining (Fig. 1A). To evaluate the relevance of these findings to humans, we used human CD8^+^ T cells derived from healthy peripheral blood mononuclear cells (PBMCs). MMA treatment caused a significant decrease in cell proliferation without changes in viability in human CD8^+^ T cells, similarly to what was observed in the murine model (Fig. 1B, Supplementary Fig. 1B).

**Fig. 1:**
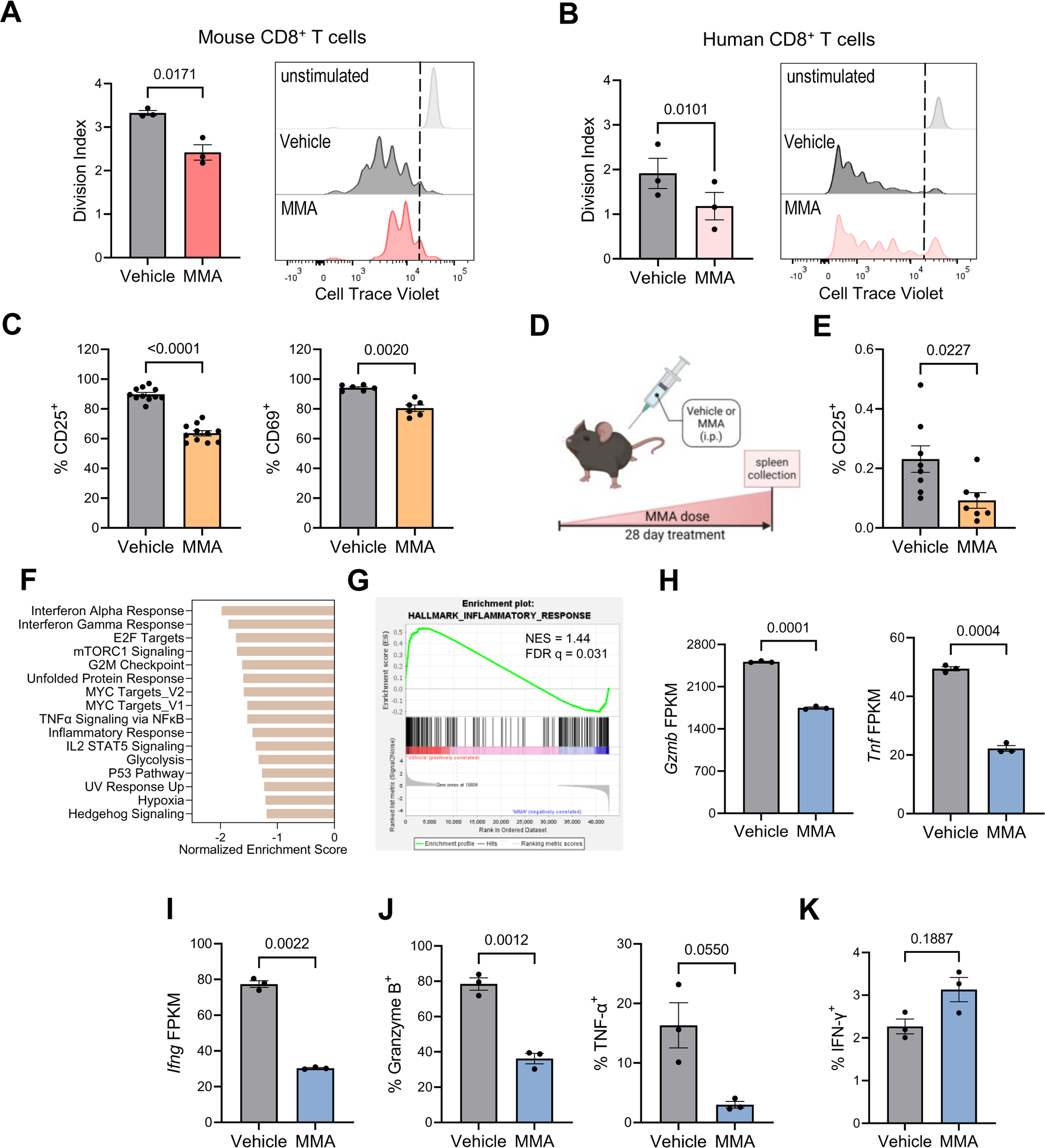
MMA impairs activation and effector functions of primary CD8^+^ T cells. **A, B** Division index (left) and representative histogram of cell trace violet staining (right) for (**A**) mouse (n = 3) and (**B**) human CD8^+^ T cells treated ± 5 mM MMA for 3 days (n = 3). The dashed line distinguishes unstimulated vs. stimulated (with anti-CD3/CD28) cells. **C** Percentage of CD25^+^ (n = 11; left) and CD69^+^ (n = 6; right) mouse CD8^+^ T cells treated ± 5 mM MMA for 3 days. **D** Schematic representation of the experimental design for administration of MMA to non-tumor-bearing C57BL/6 mice for 28 days (i.p., intraperitoneal). **E** Percentage of CD25^+^ splenic CD8^+^ T cells collected from mice administered with MMA as shown in Fig. 1D (n = 8; unpaired t-test). **F** Gene set enrichment analysis (GSEA) of hallmark transcripts statistically downregulated (false discovery rate ≤ 25%) in mouse CD8^+^ T cells treated ± 5 mM MMA for 3 days (n = 3). **G** Enrichment plot for hallmark inflammatory response (from Fig. 1F). **H** FPKM values evaluated by RNA-sequencing for *Gzmb* (granzyme B; left) and *Tnf* (TNF-α; right) of mouse CD8^+^ T cells treated ± 5 mM MMA for 3 days (n = 3). **I** FPKM values evaluated by RNA-sequencing for *Ifng* (IFN-γ) in mouse CD8^+^ T cells treated ± 5 mM MMA for 3 days (n = 3). **J** Percentage of granzyme B (left) and TNF-α (right) positive mouse CD8^+^ T cells treated ± 5 mM MMA for 3 days (n = 3). **K** Percentage of IFN-γ positive mouse CD8^+^ T cells treated ± 5 mM MMA for 3 days (n = 3). Data are represented as the mean ± SEM with statistical significance measured by paired t-tests unless otherwise indicated. Each dot represents a biological replicate.

Reductions in proliferation provoked by MMA were also associated with mouse CD8^+^ T cell activation status, as judged by significant decreases in expression of CD25 and CD69 (Fig. 1C), which are classical CD8^+^ T cell activation markers [26]. To assess the physiological relevance of these observations, we implemented a previously established protocol to raise circulatory levels of MMA in mice [21], via intraperitoneal (i.p.) administration of MMA twice daily for four weeks (Fig. 1D and Supplementary Fig. 1C). Mouse weights were recorded every other day to monitor potential toxicity of this regimen and no changes in body weight were observed between the MMA and control group (Supplementary Fig. 1D). Notably, in accord with our *in vitro* findings, assessment of splenic CD8^+^ T cells revealed a significant decrease in the expression of CD25 in the MMA-treated group (Fig. 1E).

To better understand how MMA promotes these phenotypic changes, we performed global gene expression analysis by RNA-sequencing (RNA-seq) on mouse CD8^+^ T cells treated with MMA for three days. Exposure to MMA led to the differential expression of 859 (397 downregulated and 462 upregulated) genes (Supplementary Fig. 2A). In line with phenotype analyses (Fig. 1A-1C, 1E), gene set enrichment analysis (GSEA) of the differentially expressed genes showed that MMA significantly downregulated (false discovery rate ≤ 25%) many hallmark programs associated with CD8^+^ T cell proliferation and activation, including cell cycle-related programs (e.g., E2F, MYC, p53; Fig. 1F) and metabolism (e.g., glycolysis, mTORC1 signaling; Fig. 1F).

CD8^+^ T cell activation initiates a program that drives production of cytokines and cytolytic molecules such as granzyme B (GZMB), tumor necrosis factor alpha (TNF-α), and interferon-gamma (IFN-γ) [12, 27]. GSEA analysis also indicated that exposure to MMA affected CD8^+^ T cell effector function, where there was downregulation of several inflammatory programs (e.g., IFN-α and IFN-γ responses, inflammatory processes, TNF-α signaling via NF-kB; Fig. 1F, 1G). In line with these results, RNA-seq revealed that mRNA levels of *Gzmb*, *Tnf*, and *Ifng* were decreased in CD8^+^ T cells upon exposure to MMA (Fig. 1H, 1I), which we validated by RT-qPCR (Supplementary Fig. 1E). Consistently, treatment with MMA suppressed the intracellular protein levels of granzyme B and TNF-α (Fig. 1J). Interestingly, while GSEA indicated a loss in IFN-γ response and *Ifng* mRNA levels were decreased, IFN-γ protein levels were unchanged (Fig. 1K). Thus, accumulation of MMA in circulation is sufficient to suppress CD8^+^ T cell activation and effector function, suggesting MMA is a potent immunosuppressor.

### MMA-treated CD8^+^ T cells exhibit TCA cycle abnormalities and defective oxidative phosphorylation

Metabolic deregulation is closely associated with disruption of CD8^+^ T cell activation and effector function [28, 29]. Considering the deregulation of metabolic programs upon MMA exposure observed in the GSEA and the known effects of MMA in metabolism [30], we hypothesized that MMA-induced CD8^+^ T cell dysfunction is associated with alterations of metabolic pathways that empower activation and effector function. Metabolomic analysis showed significant changes in metabolites impacted by MMA (Fig. 2A, Supplementary Fig. 2B), including deregulation of major metabolic pathways such as the pentose phosphate pathway (PPP) and the TCA cycle (Fig. 2B). Amongst these, we focused on the TCA cycle as this was the most enriched metabolic pathway and MMA is a byproduct of the propionate pathway that yields succinyl-CoA to fuel the TCA cycle (Fig. 2B, 2C). Interestingly, while oxaloacetate, citrate, and α-ketoglutarate levels remained unchanged (Supplementary Fig. 2C-2E), MMA exposure increased succinate levels and decreased fumarate levels (Fig. 2D, 2E). This is accord with previous studies showing that MMA can competitively inhibit the activity of succinate dehydrogenase (SDH), the enzyme that converts succinate to fumarate and reduces FAD^2+^ in the process [31–33], thereby leading to the accumulation of succinate and depletion of fumarate. Interestingly, malate and isocitrate, which are important metabolites for the regeneration of NADH and the proper electron flow that powers oxidative phosphorylation and ATP generation, were also decreased by MMA treatment (Fig. 2F, 2G).

**Fig. 2:**
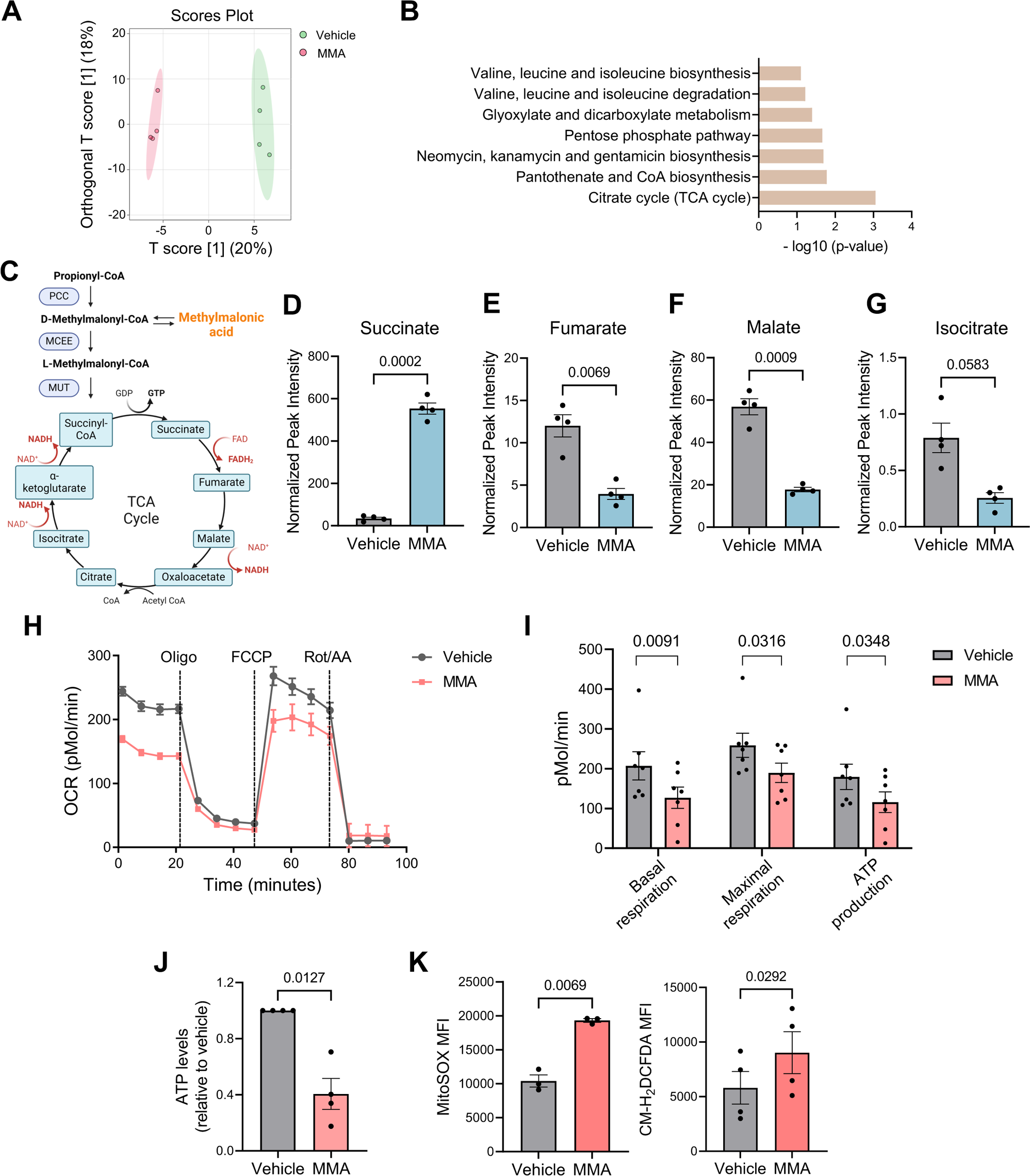
MMA-treated CD8^+^ T cells exhibit TCA cycle abnormalities and defective oxidative phosphorylation. **A** Principal component analysis (PCA) plot evaluated by metabolomic analysis of mouse CD8^+^ T cells treated ± 5 mM MMA for 3 days. **B** Pathway enrichment results for metabolic pathways statistically altered (p-value < 0.1) in mouse CD8^+^ T cells treated ± 5 mM MMA for 3 days. **C** Schematic representation of propionate pathway with connection to tricarboxylic acid (TCA) cycle. **D-G** Normalized peak intensity, as measured by metabolomic analysis of mouse CD8^+^ T cells treated ± 5 mM MMA for 3 days for the indicated metabolites: succinate, fumarate, malate, isocitrate (n = 4). **H** Oxygen consumption rate (OCR) of mouse CD8^+^ T cells treated ± 5 mM MMA for 3 days (n = 7). Dotted lines indicate addition of oligomycin (Oligo) to inhibit Complex V, carbonyl cyanide-4-(trifluoromethoxy) phenylhydrazone (FCCP) to uncouple the electron transport chain, and rotenone/antimycin A (Rot/AA) to inhibit Complex I and II, respectively. **I** Basal respiration, maximal respiration, and ATP production rate of mouse CD8^+^ T cells treated ± 5 mM MMA for 3 days (n = 7; multiple paired t-tests). **J** ATP levels of mouse CD8^+^ T cells treated ± 5 mM MMA for 3 days (n = 4). **K** MFI of mitochondrial ROS (n = 3; left) and oxidative stress (n = 4; right) of mouse CD8^+^ T cells treated ± 5 mM MMA for 3 days. Data are represented as the mean ± SEM with statistical significance measured by paired t-tests unless otherwise indicated. Each dot represents a biological replicate. Abbreviations for Fig. 2C: PCC, propionyl-CoA carboxylase; MCEE, methylmalonyl-CoA epimerase; MUT, methylmalonyl-CoA mutase; GDP, guanosine diphosphate; GTP, guanosine triphosphate; NAD, nicotinamide adenine dinucleotide; NADH, nicotinamide adenine dinucleotide hydrogen; FAD, flavin adenine dinucleotide; FADH, flavin adenine dinucleotide hydride; CoA, coenzyme A; TCA cycle, tricarboxylic acid cycle.

Given these findings the effects of MMA on mitochondrial function were assessed. Seahorse flux analyses revealed a reduction in oxygen consumption rate (Fig. 2H), including significant decreases in basal and maximal respiration, and in mitochondrial ATP production rates (Fig. 2I). These data is consistent with impairment of the electron transport chain (ETC) and oxidative phosphorylation and is in line with the decrease in reducing equivalents necessary to initiate the electron transport chain by both complex I (NADH) and complex II (FADH2) imposed by MMA treatment. In accord with these analyses, ATP levels were markedly decreased (Fig. 2J) while mitochondrial ROS, as well as total ROS levels, were increased by MMA treatment (Fig. 2K).

Thus, exposure to MMA evokes a remodeling of the TCA cycle in CD8^+^ T cells that suppresses the generation of reducing equivalents necessary to power the oxidative phosphorylation for sustained CD8^+^ T cell activation and effector function.

### MMA triggers a state of T cell exhaustion associated with the induction of TOX

Progressive loss of effector function coupled with a decrease in activation and metabolic deregulation are hallmarks of T cell exhaustion, a dysfunctional state that impairs the ability to respond to infections and/or cancer [25]. Exhausted T cells are characterized by the expression of multiple inhibitory receptors on their cell surface including PD-1, CD38, TIM-3 and LAG-3 [25, 34]. RNA-seq analyses revealed that while MMA exposure triggered the upregulation of *Pdcd1* (PD-1) and *Cd38* (CD38) mRNA levels (Fig. 3A), *Havcr2* (TIM-3) and *Lag3* (LAG-3) mRNA levels were not affected (Fig. 3B). Evaluation of the cell surface levels of these markers was consistent with these findings, where there was increased expression of PD-1 and CD38 in MMA treated CD8^+^ T cells (Fig. 3C), while TIM-3 and LAG-3 levels were unchanged (Fig. 3D).

**Fig. 3:**
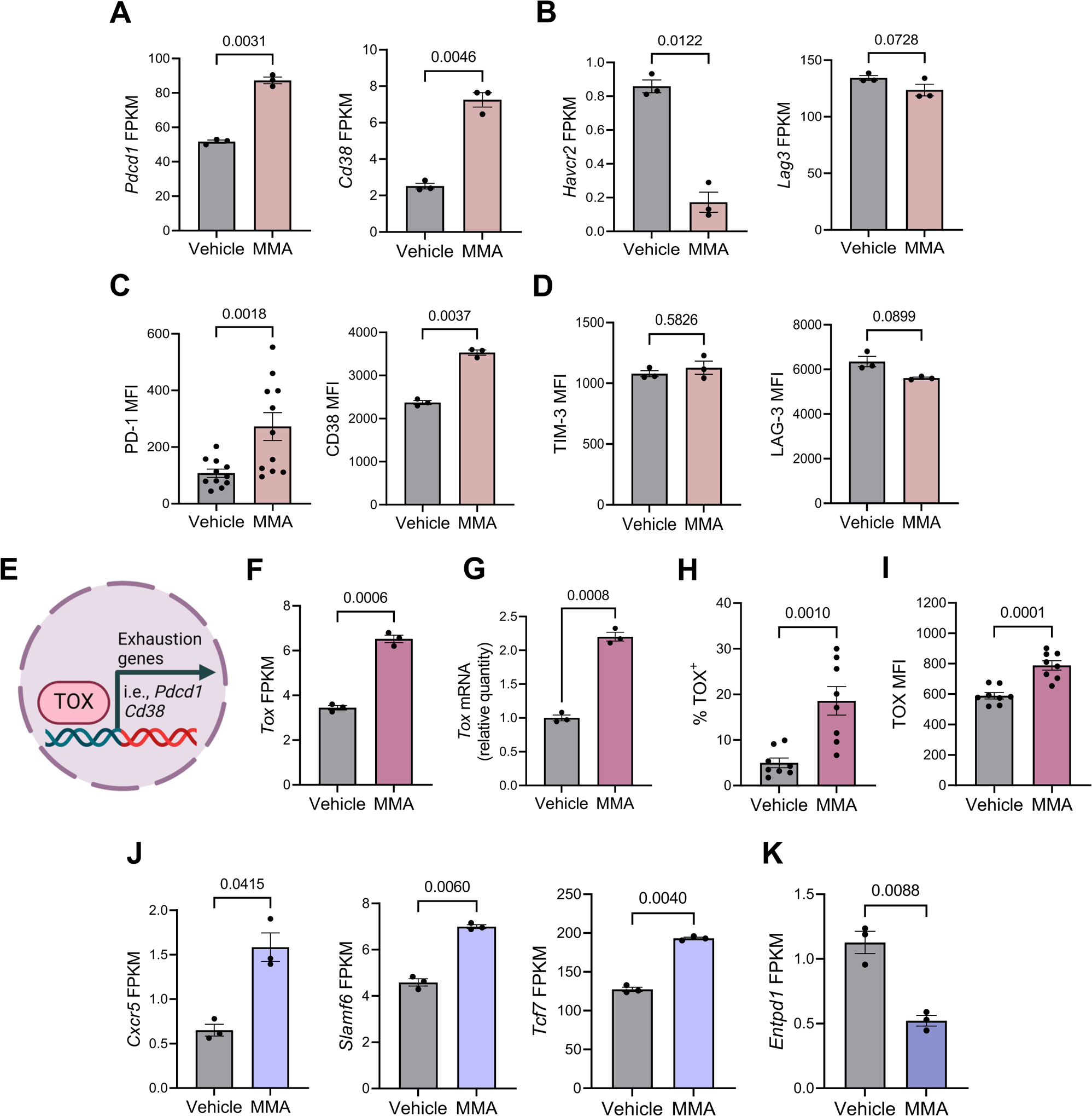
MMA induces a state of T cell exhaustion and the induction of TOX. **A** FPKM values evaluated by RNA-sequencing for *Pdcd1* (PD-1; left) and *Cd38* (CD38; right) of mouse CD8^+^ T cells treated ± 5 mM MMA for 3 days (n = 3). **B** FPKM values evaluated by RNA-sequencing for *Havcr2* (TIM-3; left) and *Lag3* (LAG-3; right) of mouse CD8^+^ T cells treated ± 5 mM MMA for 3 days (n = 3). **C** MFI of PD-1 (n = 11; left) and CD38 (n = 3; right) in mouse CD8^+^ T cells treated ± 5 mM MMA for 3 days. **D** MFI of TIM-3 (left) and LAG-3 (right) in mouse CD8^+^ T cells treated ± 5 mM MMA for 3 days (n = 3). **E** Schematic representation of the proposed model in which MMA induces TOX upregulation to promote exhaustion gene expression in CD8^+^ T cells. **F** FPKM values evaluated by RNA-sequencing for *Tox* of mouse CD8^+^ T cells treated ± 5 mM MMA for 3 days (n = 3). **G** *Tox* mRNA levels evaluated by RT-qPCR analysis of mouse CD8^+^ T cells treated ± 5 mM MMA for 3 days (n = 3). **H** Percentage of TOX^+^ splenic CD8^+^ T cells collected from mice administered MMA as shown in Fig. 1D (n = 8; unpaired t-test). **I** MFI of TOX in splenic CD8^+^ T cells collected from mice administered MMA as shown in Fig. 1D (n = 8; unpaired t-test). **J** FPKM values evaluated by RNA-sequencing for *Cxcr5* (CXCR5; left), *Slamf6* (SLAMF6; middle), and *Tcf7* (TCF-1) of mouse CD8^+^ T cells treated ± 5 mM MMA for 3 days (n = 3). **K** FPKM values evaluated by RNA-sequencing for *Entpd1* (CD39) of mouse CD8^+^ T cells treated ± 5 mM MMA for 3 days (n = 3). Data are represented as the mean ± SEM with statistical significance measured by paired t-tests unless otherwise indicated. Each dot represents a biological replicate.

Thymocyte selection-associated high mobility group box (TOX) is a transcription factor that functions as a master regulator of T cell exhaustion that induces the expression of inhibitory receptors, including PD-1 and CD38 [35–37] (Fig. 3E). Thus, we assessed the effects of MMA on TOX regulation in CD8+ T cells. Notably, *Tox* was one of the upregulated genes identified in the transcriptomic analysis, and this finding was validated by RT-qPCR (Fig. 3F, 3G). Supporting a role for TOX as the driver of CD8^+^ T cell exhaustion upon exposure to MMA, elevated MMA in circulation increased both the percentage of TOX^+^ T cells (Fig. 3H) as well as increased TOX mRNA (Fig. 3I).

CD8^+^ T cell exhaustion is a dynamic process, from activation to progenitor exhaustion through to terminal exhaustion [24, 38, 39]. As our data indicated a differential regulation of the exhaustion driven inhibitory markers, markers that distinguish these subsets of exhausted T cells were analyzed using the RNA-seq dataset. Interestingly, MMA exposure increased mRNA levels of markers of progenitor exhaustion, including *Cxcr5* (CXCR5), *Slamf6* (SLAMF6), and *Tcf7* (TCF-1) (Fig. 3J). On the other hand, MMA exposure decreased mRNA levels of the terminal exhaustion marker, *Entpd1* (CD39; Fig. 3K), in accord with the progenitor exhaustion phenotype [40, 41]. Thus, exposure to MMA triggers the induction of TOX, which drives the induction of an early exhaustion program in CD8^+^ T cells.

### MMA reduced CD8^+^ T cell cytotoxicity and increased tumor penetrance

Given these findings we reasoned that exposure to MMA might impair the cytotoxic activity of CD8^+^ T cells against tumor cells. In support of this notion, treatment of lung tumor cells derived from KP mice (genetically engineered LSL-Kras^G12D^;Trp53^flox/flox^ lung cancer mouse model [22]) with conditioned media from CD8^+^ T cells that were pre-treated with MMA showed reduced cytotoxicity compared to KP cells treated with conditioned media from CD8^+^ T cells from control conditions (Supplementary Fig. 3A and 3B). To validate these observations, CD8^+^ T cells were isolated from OT-I mice, whose T cell receptor (TCR) is engineered to specifically respond to the class I-restricted ovalbumin antigen (OVA 257-264) [42], and treated them with MMA upon activation for three days. Activated CD8^+^ OT-I cells were then co-cultured with OVA-expressing Lewis Lung Carcinoma cells (LLC-OVA; Fig. 4A). As expected, CD8^+^ OT-I cells induced death of LLC-OVA, yet MMA treatment significantly reduced in CD8^+^ OT-I-induced cell death (Fig. 4B); thus, MMA impairs CD8^+^ T cell anti-tumor cytotoxicity. Finally, to assess how elevated MMA levels in circulation affects tumorigenesis *in vivo*, we challenged control and MMA-treated mice with orthotopically transplanted KP tumor cells expressing luciferase, a powerful immunogen in C57BL/6 mice [43], (KP-Luciferase cells) (Fig. 4C). Systemic levels of MMA were elevated by the administration of MMA even in the presence of tumors (Fig. 4D). Importantly, while the lungs of the control group only had 40% tumor penetrance (2/5 mice; Fig. 4E, 4F), mice exposed to elevated levels of MMA in circulation displayed 100% tumor penetrance (6/6 mice; Fig. 4E, 4F). Thus, exposure to MMA impairs CD8^+^ T cell cytotoxic activity towards lung cancer cells, enabling tumor progression.

**Fig. 4:**
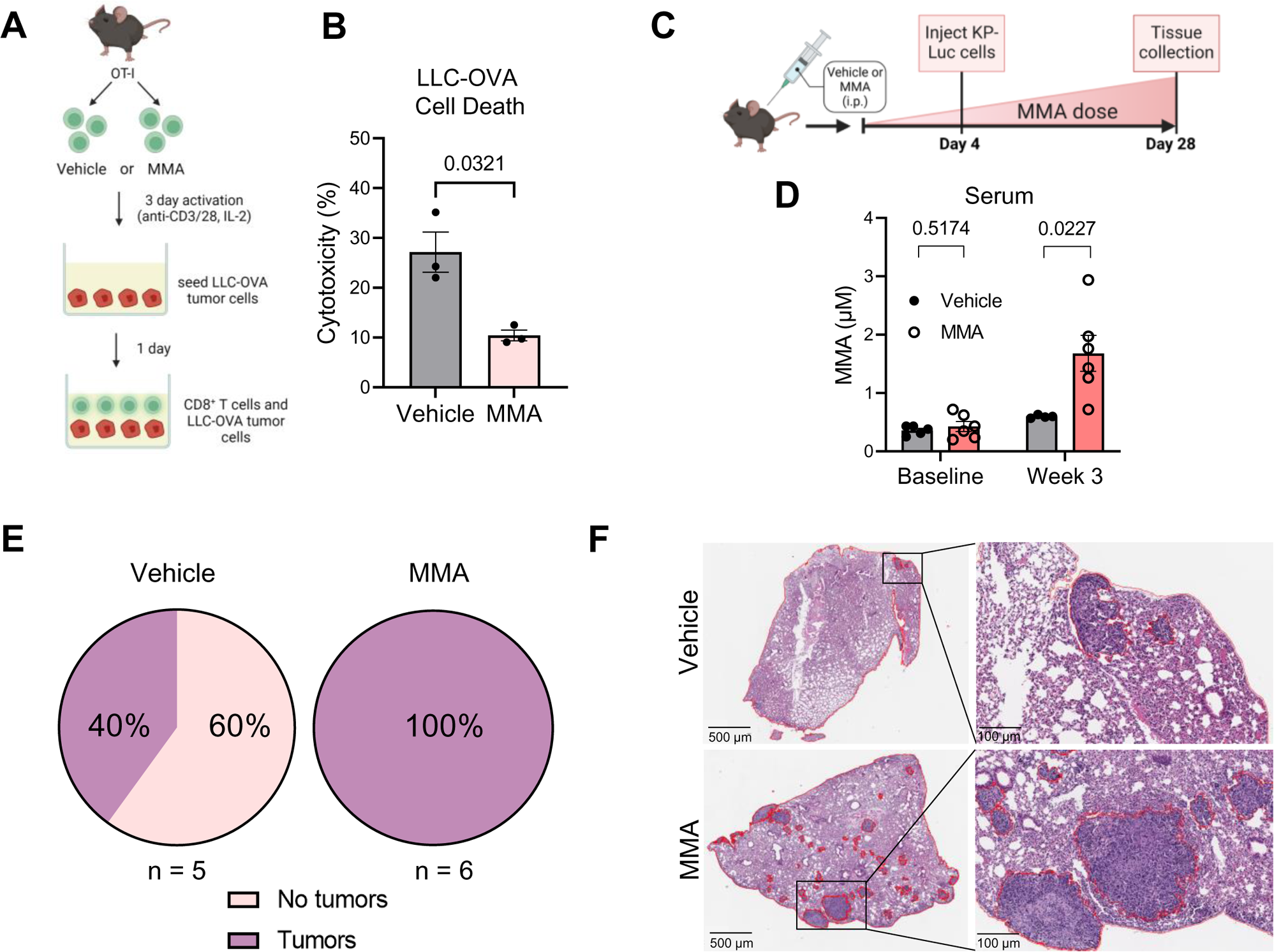
MMA impairs CD8^+^ T cell cytotoxicity and augments tumor penetrance. **A** Schematic representation of the experimental design to assess cytotoxicity of CD8^+^ OT-I cells treated ± 5 mM MMA against OVA-expressing LLC cells (LLC-OVA). **B** Cytotoxicity, as measured by percentage of LLC-OVA cell death after 24 hour co-culture with CD8^+^ OT-I cells treated ± 5 mM MMA for 3 days (n = 3). **C** Schematic representation of the experimental design for administration of MMA to KP-Luciferase (KP-Luc) tumor-bearing C57BL/6 mice for 28 days (i.p., intraperitoneal). **D** MMA concentration in serum at indicated time points collected from mice administered MMA as shown in Fig. 4C (n = 5 or 6 per group; multiple unpaired t-tests). **E** Representation of tumor penetrance, as measured by the presence of at least one tumor in H&E lung slides. **F** Representative H&E slides at 500 μm (left) and 100 μm (right) of lungs collected from tumor-bearing mice administered MMA as shown in Fig. 4C. Data are represented as the mean ± SEM with statistical significance measured by paired t-tests unless otherwise indicated. Each dot represents a biological replicate.

## Discussion

Circulatory MMA levels increase with age [3–6] and recent work has revealed MMA can promote cancer progression [6, 9, 44], suggesting MMA as an important link between aging and cancer. Here, we add a new dimension to the contribution of MMA to cancer, where we show that that exposure to MMA triggers exhaustion of activated CD8^+^ T cells, thereby halting anti-tumor immunity and enabling the manifestation of malignant phenotypes. Mechanistically, we ascribe this effect to the metabolic defects imposed by MMA and to the induction of the exhaustion master regulator, TOX. While previous work has shown that TCA cycle deregulation and suppression of oxidative phosphorylation can provoke T cell exhaustion [45, 46], additional studies are needed to confirm whether these metabolic alterations are the root cause of the observed T cell exhaustion.

Our analysis of different exhaustion markers suggest that, at least under the tested exposure paradigm, MMA triggers an early state of exhaustion designated as “progenitor exhaustion” [47, 48], with the absence of markers of terminally exhausted T cells (TIM-3^+^TCF-1^−^ [38]). Moreover, our data show that MMA impairs CD8^+^ T cell effector function by suppressing the production of granzyme B and TNF-α but not IFN-γ. In a model of chronic infection, loss of CD8^+^ T cell effector function has been shown to occur in a hierarchical manner, where loss of IFN-γ production is more resistant to exhaustion and occurs later compared to loss of TNF-α production [49]. Importantly, while more work is needed to precisely define the dynamics of MMA-induced CD8^+^ T cell exhaustion and its clinical significance, it has been shown that in patients with T cell/histiocyte-rich large B-cell lymphoma (THRLBCL), TCF1^+^ progenitor exhausted T cells correlate with good clinical response to anti-PD-1 immunotherapy, while terminally exhausted T cells do not [50]. Thus, circulatory levels of MMA might constitute an important and simple predictive biomarker of immunotherapy response.

Our findings add MMA to the growing body of metabolites with immunomodulatory roles and lays the foundation for MMA as a link between whole body physiology (e.g., aging, vitamin B12 deficiency), immune deregulation, and the development of pathologies like cancer. Additional studies aimed at defining the mechanisms by which MMA exerts its immunomodulatory roles in CD8^+^ T cells, as well as its potential effects on other immune cell populations and their concerted actions in the TME, will provide new insights into the full spectrum of the effect of MMA on tumorigenesis.

## Acknowledgements

We would like to thank members of the DeNicola lab (Moffitt) for helpful feedback and support; Dr. Brian Ruffell and members of the Ruffell lab (Moffitt) for kindly sharing LLC-ZsGreen and LLC-OVA-ZsGreen cells as well as helpful advice; Dr. Dorina Avram (Moffitt) for kindly providing the OT-I mice; members of Dr. Javier Pinilla-Ibarz lab (Moffitt) for kindly sharing human PBMCs; and Drs. Alfred Zippelius (University of Basel) and Massimo Broggini (Instituto di Ricerche Farmacologiche Mario Negri IRCCS) for kindly sharing KP1.9 cells. We would also like to thank the Moffitt Cancer Center/USF Comparative Medicine Program for animal care; staff members of the Flow Cytometry, Proteomics and Metabolomics, and Analytic Microscopy core facilities at the H. Lee Moffitt Cancer Center & Research Institute, an NCI designated Comprehensive Cancer Center (P30-CA076292). RSH is supported by the T32 CA233399 program. JLC is supported by the Cortner-Couch Endowed Chair for Cancer Research from the University of South Florida School of Medicine and by P01-CA250984. The mass spectrometry work was partially funded by NIH grants 5P01CA120964 (JMA) and 5P30CA006516 (JMA). APG is supported by the Pathway to Independence Award from NCI (R00CA218686; R00CA218686-04S1), the New Innovator Award from OD/NIH (DP2AG0776980), an American Cancer Society Research Scholar Award (RSG-22-164-01-MM), the NIA (R21AG083720), METAvivor, the Florida Breast Cancer Research Foundation, the Phi Beta Psi Sorority, the American Lung Association, and the Florida Health Department Bankhead-Coley Research Program. Schematic figures were created using BioRender.com.

## Author Contributions

JDT contributed to experiment design and execution, data analysis, preparation of figures, and manuscript writing. RSH and SD contributed to experiment design and execution, data analysis, and interpretation. DI prepared the RNA for the RNA-seq experiments and performed its analysis. SD, DR, FL, and HK contributed to tissue collection and processing of mouse experiments. JF-G and S-MF contributed to methodology and identification of MMA-regulated metabolism. ML performed the mass spectrometry analysis of MMA in mouse serum. JLC contributed to project supervision, interpretation of results, and provided critical reagents. JA performed the metabolomic analysis. APG contributed to project conception and design, supervision, data analysis, data interpretation, funding acquisition, and manuscript writing. All authors have reviewed and approved the manuscript.

## Data Availability Statement

Data will be made available upon reasonable request to

## Supplementary Information

Are provided in a separate Word document titled “Supplementary Information”.

## Subject Ontology

Tumor immunology; Cancer metabolism

