## Supplemental Information for "Methylmalonic acid induces metabolic abnormalities and exhaustion in CD8^+^ T cells to suppress anti-tumor immunity"

[1]Department of Molecular Medicine, University of South Florida Morsani College of Medicine, Tampa, FL, 33612, USA
[2]Department of Molecular Oncology, H. Lee Moffitt Cancer Center & Research Institute, Tampa, FL, 33612, USA
[3]Department of Tumor Microenvironment and Metastasis, H. Lee Moffitt Cancer Center & Research Institute, Tampa, FL, 33612, USA
[4]Proteomics and Metabolomics Core, H. Lee Moffitt Cancer Center & Research Institute, Tampa, Florida, 33612, USA
[5]Laboratory of Cellular Metabolism and Metabolic Regulation, VIB-KU Leuven Center for Cancer Biology, VIB, 3001 Leuven, Belgium
[6]Department of Medicine, Beth Israel Deaconess Medical Center and Harvard Medical School, Boston, MA, 02215, USA

Contents:

Supplementary Methods

Supplementary Table 3: Key Materials

Supplementary Table 4: Primers for RT-qPCR analysis

Supplementary Figures and Figure Legends (1-3)

Supplementary References

### Supplementary Methods

#### Human $CD8^+$ T cell isolation and culture conditions

Healthy human donor peripheral blood mononuclear cells (a gift from Dr. Pinilla-Ibarz) (n = 3; 35 year old male, 21 year old female, 20 year old female) were derived from buffy coats (Life South Community Blood Bank, Brooksville, FL, USA) using Ficoll-Paque PLUS density gradient centrifugation (Cytiva) and frozen at -80°C in 90% FBS and 10% DMSO prior to use. Human $CD8^+$ T cells were isolated using negative immunomagnetic selection (Miltenyi Biotec, Bergisch Gladbach, Germany, ~80% purity) with LS Columns (Miltenyi Biotec) and were cultured in RPMI 1640 media (Cytiva) supplemented with 10% FBS (Gibco) and 1% penicillin/streptomycin (Cytiva). Human T cells were rested in media overnight at 37°C and 5% $CO_2$ before activation. Cells were seeded at a concentration of 200,000 cells/well in 96-well round-bottom plates that were coated overnight with 5 μg/mL of anti-human CD3ε antibody (BioLegend) and washed twice with PBS prior to seeding. T cell activation was performed using 1 μg/mL anti-human CD28 antibody (Thermo Fisher Scientific) in the presence of 10 ng/mL recombinant human IL-2 (PeproTech, Cranbury, NJ, USA); additionally, MMA (Sigma-Aldrich), solubilized in molecular grade water, was added at a concentration of 5 mM for up to 5 days at 37°C and 5% $CO_2$.

#### Cell lines

KP-Luciferase [1] (a gift from Dr. Zippelius), LLC-ZsGreen [2] (a gift from Dr. Ruffell), LLC-OVA-ZsGreen [2] (a gift from Dr. Ruffell) cell lines were cultured in DMEM high glucose medium (Cytiva) with 10% FBS (Gibco) and 1% penicillin/streptomycin (Cytiva) at 37°C and 5% $CO_2$. All cell lines tested negative for mycoplasma.

#### Mitochondrial function assays

After three days of activation in the presence of 5 mM MMA, mouse $CD8^+$ T cells were harvested, washed with Seahorse XF media (Agilent, Santa Clara, CA, USA), and seeded at a concentration of 200,000 cells/well in a 96-well Seahorse culture plate (Agilent) prior to non-$CO_2$ incubation for 60 min. Seahorse XFe96 Metabolic Extracellular Flux Analyzer (Agilent) was used, and the XF Cell Mito Stress Test assay was run according to manufacturer's instructions. Cells were stimulated with oligomycin (1 μM), FCCP (1.5 μM), and rotenone/antimycin A (0.5 μM). Oxygen consumption rates (OCR) were analyzed using Wave Software v2.4.3.3 (Agilent).

**ATP assay**

After three days of activation in the presence of 5 mM MMA, mouse $CD8^+$ T cells were harvested, and ATP levels were assessed using a luminescence-based ATP Detection Assay Kit (Cayman Chemical Company, Ann Arbor, MI, USA) according to manufacturer's instructions.

**Histology (H&E) imaging**

Lung tissues were fixed with 10% formalin overnight, transferred to 70% ethanol, and paraffin embedded to be sectioned. Haematoxylin and eosin (H&E) staining was performed by IDEXX BioAnalytics (Westbrook, ME, USA). Slides were scanned with an Aperio imager (Leica Biosystems, Nussloch, Germany) at 20x and analyzed for the presence of tumors using QuPath v.0.3.0 software [3].

**Targeted metabolomics**

Metabolites were extracted using 80% (v/v) aqueous methanol from mouse $CD8^+$ T cells treated with vehicle or 5 mM MMA upon activation for three days. Samples were re-suspended using 20 μL HPLC grade water for mass spectrometry. 5-7 μL were injected and analyzed using a hybrid 6500 QTRAP triple quadrupole mass spectrometer (AB/SCIEX, Framingham, MA, USA) coupled to a Prominence UFLC HPLC system (Shimadzu, Kyoto, Japan) via selected reaction monitoring (SRM) of a total of 300 endogenous water-soluble metabolites for steady-state analyses of samples. Some metabolites were targeted in both positive and negative ion mode for a total of 311 SRM transitions using positive/negative ion polarity switching. ESI voltage was +4950V in positive ion mode and –4500V in negative ion mode. The dwell time was 3 ms per SRM transition and the total cycle time was 1.55 seconds. Approximately 9-12 data points were acquired per detected metabolite. Samples were delivered to the mass spectrometer via hydrophilic interaction chromatography (HILIC) using a 4.6 mm i.d x 10 cm Amide XBridge column (Waters, Milford, MA, USA) at 400 µL/min. Gradients were run starting from 85% buffer B (HPLC grade acetonitrile) to 42% B from 0-5 minutes; 42% B to 0% B from 5-16minutes; 0% B was held from 16-24 minutes; 0% B to 85% B from 24-25 minutes; 85% B was held for 7 minutes to re-equilibrate the column. Buffer A was comprised of 20 mM ammonium hydroxide/20 mM ammonium acetate (pH = 9.0) in 95:5 water:acetonitrile. Peak areas from the total ion current for each metabolite SRM transition were integrated using MultiQuant v3.0.2 software (AB/SCIEX). Statistical analysis was performed using MetaboAnalyst v4.0, a free online software for the analysis of metabolomic experiments (https://www.metaboanalyst.ca/). Normalization of the original data was done using the median of the entire metabolome for each sample and log-transformed before further analysis. Data are provided in Supplementary Table 1.

**Quantification of MMA concentration in mouse serum**

Stable isotope-labelled MMA ($^{13}C_4$, 99%; Cambridge Isotope Labs, Tewksbury, MA, USA) was added to 50 µL of mouse serum, added to a 96 well plate, and diluted by adding 250 µL of water per well. Oasis MAX µElution plate (Waters) were conditioned with 250 µL 100% methanol followed by 300 µL water. After each condition, positive pressure was applied to the plate via Waters Positive Pressure-96 Processor. The diluted serum samples were added to the µElution plate and washed with 300 µL of 5% ammonium hydroxide (Sigma-Aldrich) followed by 250 µL of 100% acetonitrile (Burdick and Jackson, Honeywell, Muskegon, MI, USA). Metabolites, including MMA, were eluted from the µElution plate by adding 200 µL of 2% Formic acid in methanol and dried by blowing nitrogen gas stream. The metabolites were re-dissolved in 80% acetonitrile and 20% of 10 mM ammonium carbonate/ammonium hydroxide (Sigma-Aldrich). Ultra-high performance liquid chromatography-high resolution mass spectrometry (UHPLC-HRMS) was performed using a Vanquish UHPLC interfaced with a Q Exactive FOCUS quadrupole-orbital ion trap mass spectrometer (Thermo Fisher Scientific). Chromatographic separation was performed using an Atlantis Premier BEH Z-HILIC Column (2.1 mm ID x 100 mm length, 1.7 µm particle size). Mobile phase A was aqueous 10 mM ammonium carbonate and 0.05% ammonium hydroxide, and mobile phase B was 100% acetonitrile. The gradient program included the following steps: start and stay at 80% B for 2 minutes, a linear gradient from 80 to 20% B over 10 minutes, stay at 20% B for 0.5 minutes, return to 80% B in 0.5 minutes, and re-equilibration for 2 minutes for a total run time of 15 minutes. The flow rate was set to 0.25 mL/min. The autosampler was cooled to 5°C and the column temperature was set to 30°C. Sample injection volume was 5 µL. Full MS at negative polarity was used for mass spectrum data acquisition. Data analysis was performed using Xcalibur v4.5 (Thermo Fisher Scientific).

**Quantitative Real-Time PCR**

After three days of activation in the presence of 5 mM MMA, mouse $CD8^+$ T cells were harvested, and RNA was isolated using RNeasy Mini Kit (Qiagen, Hilden, Germany) and contaminant DNA was digested on column with RNase-free DNase (Qiagen). cDNA was synthesized using the iSCRIPT cDNA synthesis kit (BioRad, Hercules, CA, USA) and analyzed on a QuantStudio 6 Pro Real-Time PCR system (Thermo Fisher Scientific) using SYBR green master mix (Life Technologies, Carlsbad, CA, USA). Gene expression was analyzed using the Design and Analysis software v2.6.0 (Life Technologies) and further processed using Microsoft Excel version 2302 (Microsoft, Redmond, WA, USA). Target gene expression was normalized to expression of TATA-binding protein (TBP). A list of sequences for primers used is provided in Supplementary Table 4.

**Global gene expression analysis (RNA-Sequencing)**

Total RNA was isolated and DNase treated as described above, and then sent to Novogene (Sacramento, CA, USA) for further processing and RNA-seq analysis. Briefly, RNA quality was assessed using the 5400 Fragment Analyzer system (Agilent) and all samples had RIN score of 9.8. mRNA was purified from total RNA using poly-T oligo-attached magnetic beads and first strand cDNA was synthesized using random hexamer primers followed by second strand cDNA synthesis using dUTP for directional library or dTTP for non-directional library. Quantified libraries were pooled and sequenced on Novaseq 6000, according to effective library concentration and data amount. The library construction was created using NEB ultra kit according to manufacturer's instructions. Raw data (raw reads) of fastq format were firstly processed through Novogene in-house perl scripts. In this step, clean data (clean reads) were obtained by removing reads containing adapters, reads containing ploy-N and low quality reads from raw data. Index of the reference genome (Mus Musculus: GRCm39/mm39) was built using Hisat2 v2.0.5 and paired-end clean 1 reads were aligned to the reference genome using Hisat2 v2.0.5. featureCounts v1.5.0-p3 was used to count the reads numbers mapped to each gene and FPKM of each gene was calculated based on gene length and reads count mapped to this gene. The accession number for the raw sequencing data reported in this paper is GEO: GSE256487 and can be accessed on https://www.ncbi.nlm.nih.gov/geo/. Differential expression analysis was performed using the DESeq2 through the interactive web application (DEApp) developed in R with Shiny (https://yanli.shinyapps.io/DEApp/, developed by the bioinformatics core (Center for Research Informatics (CRI), University of Chicago). FDR adjusted p-value of 0.05 and absolute fold change of 1.5 were set as the threshold for analysis. A list of differentially expressed genes is provided in Supplementary Table 2. Gene Set Enrichment Analysis (GSEA) for mouse-ortholog hallmark gene sets (https://www.gsea-msigdb.org/gsea/index.jsp) [4-7] was performed using the GSEA software v.4.3.2 (a joint project of UC San Diego and Broad Institute) [4, 8]. Only gene sets with FDR adjusted p-value ≤ 25% were evaluated.

### Supplementary Table 3: Key Materials

| Antibody | Source | Identifier |
|---|---|---|
| Mouse CD3ε, ultra-LEAF purified | Biolegend | 100359, clone 145-2C11 |
| Mouse CD28, ultra-LEAF purified | BioLegend | 102116, clone 37.51 |
| Mouse CD3, PE/Cy7 | Tonbo | 60-0032-U100, clone 17A2 |
| Mouse CD8, BUV395 | BD Biosciences | 563786, clone 53-6.7 |
| Mouse CD25, FITC | BioLegend | 101908, clone 3C7 |
| Mouse CD69, PE/Cy7 | BioLegend | 104512, clone H1.2F3 |
| Mouse PD-1, AF647 | BioLegend | 135230, clone 29F.1A12 |
| Mouse CD38, FITC | BioLegend | 102705, clone 90 |
| Mouse Granzyme B, AF647 | Sony Biotechnology | 3177025, clone GB11 |
| Mouse TNF-α, PE/Cy7 | BioLegend | 506323, clone MP6-XT22 |
| Mouse IFN-γ, FITC | BioLegend | 505806, clone XMG1.2 |
| Mouse TOX, eFluor 660 | Thermo Fisher Scientific | 50-6502-82, clone TXRX10 |
| CD16/CD32 (Mouse Fc block) | BD Biosciences | 553142, clone 2.4G2 |
| Human CD3ε, ultra-LEAF purified | BioLegend | 317347, clone OKT3 |
| Human CD28 Monoclonal | Thermo Fisher Scientific | 16-0289-85, clone CD28.2 |
| Human CD3, BV711 | Biolegend | 300463, clone UCHT1 |
| Human CD8, FITC | Tonbo | 35-0089-T100, clone HIT8a |
| **Chemical** | **Source** | **Identifier** |
| Methylmalonic acid | Sigma-Aldrich | M54058 |
| Mouse IL-2 recombinant protein | BioLegend | 575404 |
| Human IL-2 recombinant protein | PeproTech | 200-02 |
| Ficoll-Paque PLUS | Cytiva | 17144002 |
| Cell Trace Violet | Thermo Fisher Scientific | C34571 |
| CM-$H_2$DCFDA | Thermo Fisher Scientific | C6827 |
| MitoSOX Red | Thermo Fisher Scientific | M36008 |
| Protein transport inhibitor with brefeldin A | BD Biosciences | 555029 |
| Propidium iodide | Sigma-Aldrich | P4170 |
| Fixable Near-IR viability | Thermo Fisher Scientific | L34976 |
| Zombie Yellow viability | Biolegend | 423103 |
| Sytox Deep Red | Thermo Fisher Scientific | S11381 |
| Oligomycin | Sigma-Aldrich | O4876 |
| FCCP | Sigma-Aldrich | C2920 |
| Antimycin A | Sigma-Aldrich | A8674 |
| Rotenone | Cayman Chemical Company | 13995 |
| RNase-free DNase | Qiagen | 79254 |

| | | |
|---|---|---|
| RBC lysis buffer | Biolegend | 420301 |
| EDTA | Thermo Fisher Scientific | AM9260G |
| RPMI-1640 medium | Cytiva | SH30027FS |
| DMEM high glucose medium | Cytiva | SH30081 |
| Fetal bovine serum (FBS) | Gibco | 26140079 |
| β-Mercaptoethanol | Sigma-Aldrich | M3148 |
| Sodium Pyruvate | Gibco | 11360070 |
| MEM nonessential amino acids | Gibco | 11140050 |
| HEPES | Thermo Fisher Scientific | 15630106 |
| Penicillin-streptomycin | Cytiva | SV30010 |
| **Assay/Kit** | **Source** | **Identifier** |
| MojoSort Mouse CD8$^+$ Naïve T cell Isolation Kit | BioLegend | 480044 |
| Human Pan T cell Isolation Kit | Miltenyi Biotec | 130-096-535 |
| Seahorse XF Cell Mito Stress Test Kit | Agilent | 103015-100 |
| eBioscience Foxp3/Transcription Kit | Thermo Fisher Scientific | 00-5523-00 |
| RNeasy Mini Kit | Qiagen | 74104 |
| ATP Detection Assay Kit | Cayman Chemical Company | 700410 |
| **Other** | **Source** | **Identifier** |
| 70 µm nylon cell strainer | Thermo Fisher Scientific | 22-363-548 |
| 96-well round-bottom plate | Sarstedt | 83.3925 |
| 96-well flat bottom plate | Sarstedt | 83.3924 |
| LS Columns | Miltenyi Biotec | 130-042-401 |

**Supplementary Table 4: Primers for RT-qPCR analyses**

| Gene | Sequence 5' to 3' |
|---|---|
| Mouse *Gzmb* Forward | CCTCCAGGACAAAGGCAG |
| Mouse *Gzmb* Reverse | CAGTCAGCACAAAGTCCTCTC |
| Mouse *Ifng* Forward | CCTAGCTCTGAGACAATGAACG |
| Mouse *Ifng* Reverse | TTCCACATCTATGCCACTTGAG |
| Mouse *Tox* Forward | TGCTCTCCAATTCCATCTCTG |
| Mouse *Tox* Reverse | CTGTCTGATGTCTGTAGGCTG |
| Mouse *TBP* Forward | AGAACAATCCAGACTAGCAGCA |
| Mouse *TBP* Reverse | GGGAACTTCACATCACAGCTC |

### Supplementary Figures and Figure Legends (1-3)

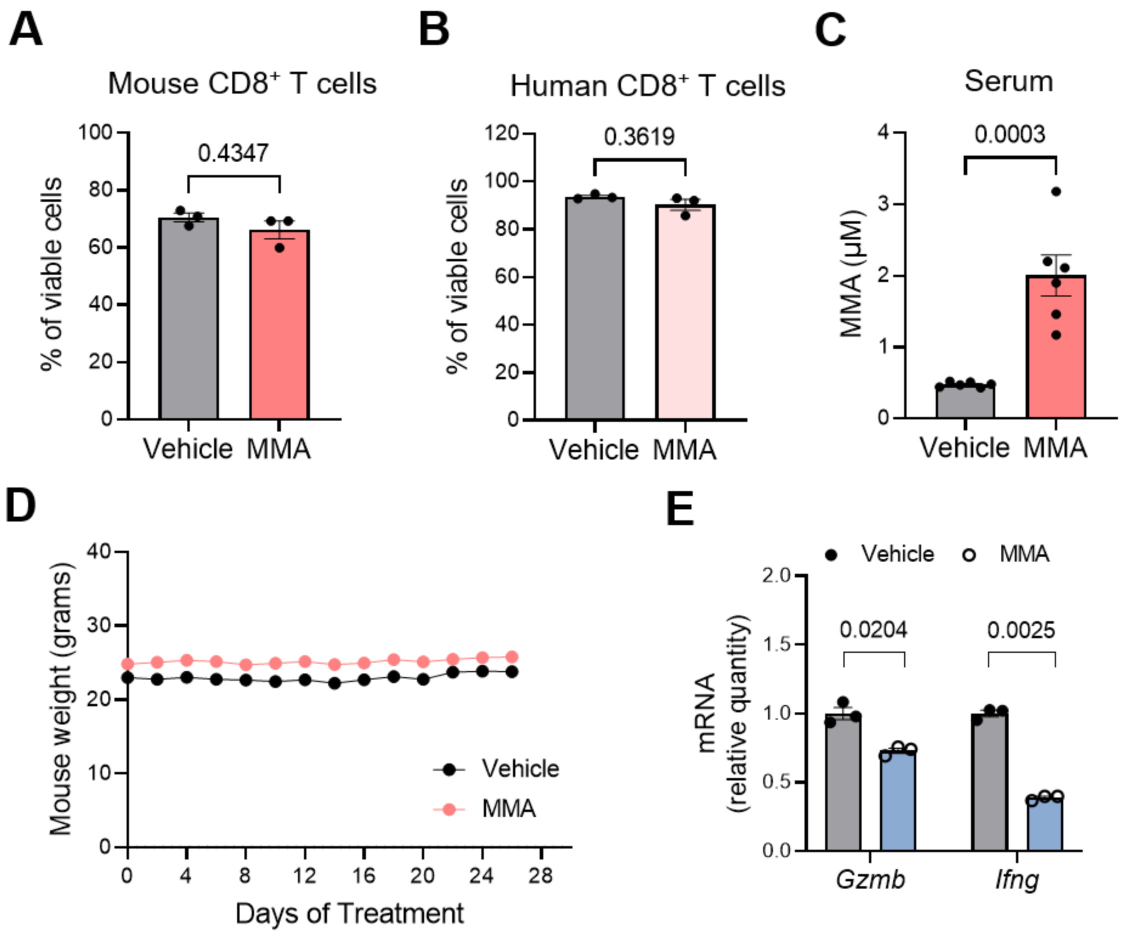


**Supplementary Fig. 1: A** Percentage of viable cells in mouse (n = 3) and **B** human CD8+ T cells treated ± 5 mM MMA for 3 days (n = 3). **C** MMA concentration in serum collected at week 3 from mice administered MMA as shown in Fig. 1D (n = 6; unpaired t-test). **D** Body weights of non-tumor-bearing C57BL/6 mice that were intraperitoneally injected with saline or MMA twice daily for 28 days as shown in Fig. 1D (n = 8). **E** *Gzmb* and *Ifng* mRNA levels evaluated by RT-qPCR analysis of mouse CD8+ T cells treated ± 5 mM MMA for 3 days (n = 3). Data are represented as the mean ± SEM with statistical significance measured by paired t-tests unless otherwise indicated. Each dot represents a biological replicate.

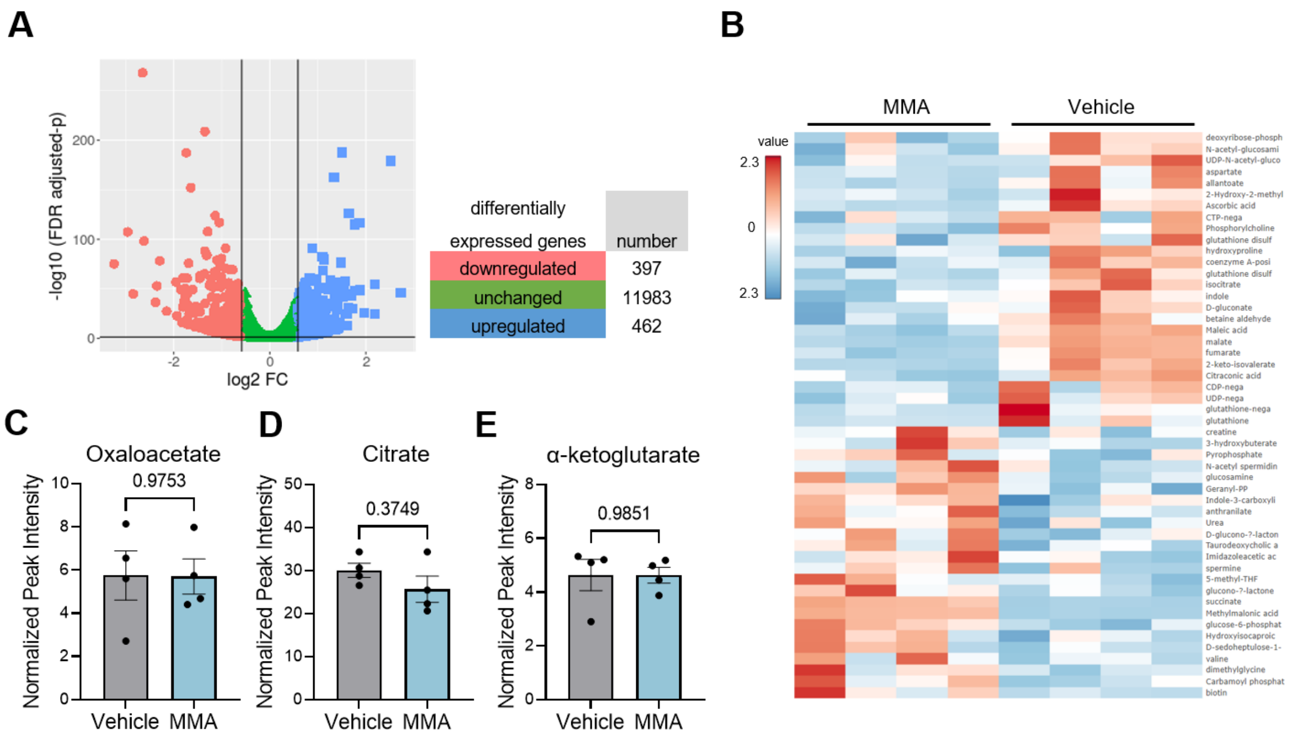


**Supplementary Fig. 2: A** Volcano plot representing transcripts that were differentially expressed (FDR adjusted p-value ≤ 0.05) in mouse CD8$^+$ T cells treated ± 5 mM MMA for 3 days (n = 3). **B** Heat map for metabolomic analysis of CD8$^+$ T cells treated ± 5 mM MMA for 3 days (n = 4). **C-E** Normalized peak intensity, as measured by metabolomic analysis of mouse CD8$^+$ T cells treated ± 5 mM MMA for 3 days for the indicated metabolites: oxaloacetate, citrate, α-ketoglutarate (n = 4). Data are represented as the mean ± SEM with statistical significance measured by paired t-tests unless otherwise indicated. Each dot represents a biological replicate.

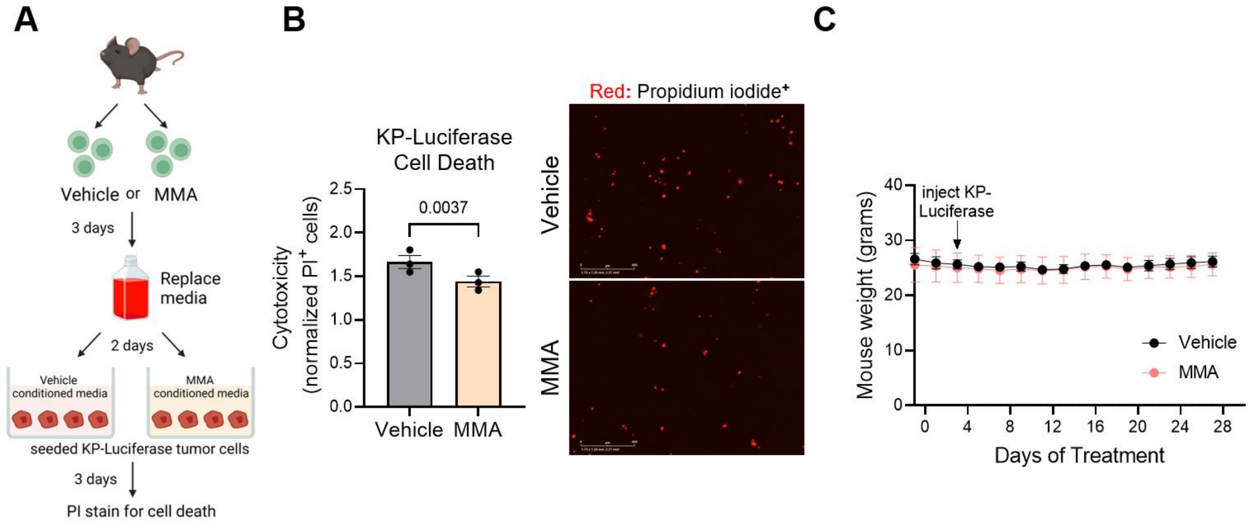


**Supplementary Fig. 3: A** Schematic representation of experimental design to assess cytotoxicity effects of vehicle or MMA-treated conditioned media on KP-Luciferase cell death. **B** Measure of cytotoxicity (n = 3; left) and representative images at 4x magnification depicting cells positive for propidium iodide (right). **C** Body weights of C57BL/6 mice (n = 5 or 6 per group) that were intraperitoneally injected with saline or MMA twice daily for 28 days. At day 4, 100,000 KP-Luciferase cells were injected via tail vein to all mice, as shown in Fig. 4C. Data are represented as the mean ± SEM with statistical significance measured by paired t-tests unless otherwise indicated. Each dot represents a biological replicate.
